# Future sea-level rise drives rocky intertidal habitat loss and benthic community change

**DOI:** 10.1101/553933

**Authors:** Nikolas J. Kaplanis, Clinton B. Edwards, Yoan Eynaud, Jennifer E. Smith

**Author notes:** Corresponding author: (NJK).

## Abstract

Rocky intertidal ecosystems may be particularly susceptible to sea-level rise impacts but few studies have explored community scale response to future sea-level scenarios. Combining remote-sensing with large-area imaging, we quantify habitat extent and describe biological community structure at two rocky intertidal study locations in California. We then estimate changes in habitat area and community composition under a range of sea-level rise scenarios using a model-based approach. Our results suggest that future sea-level rise will significantly reduce rocky intertidal area at our study locations, leading to an overall decrease in benthic habitat and a reduction in overall invertebrate abundances, but increased densities of certain taxa. These results suggest that sea-level rise may fundamentally alter the structure and function of rocky intertidal systems. As large scale environmental changes such as sea-level rise accelerate in the next century, more extensive spatially-explicit monitoring at ecologically relevant scales will be needed to visualize and quantify the impacts to biological systems.

## Introduction

### Sea-level rise projections and potential impacts to the rocky intertidal

Sea-level rise is predicted to alter habitat availability and modify community structure in many marine ecosystems [1,2]. In the monitoring record, sea-level changes vary among locations but generally show a rising trend, with rates showing a recent increase [1,3–9]. While projections remain uncertain, recent studies predict extremes of up to 2.5 meters of sea-level rise within the next century [3,10]. Such large magnitude and rapid sea-level rise poses a substantial risk to the integrity of coastal ecosystems, yet the extent to which it will modify the physical and ecological structure of rocky coastlines remains mostly unknown [11–14].

Rocky intertidal systems may be particularly vulnerable to sea-level rise driven habitat loss [11–14]. When backed by steep cliffs or anthropogenic structures, as is the case on the majority of coastlines globally [15], rather than migrating shoreward with sea-level rise, many rocky shores are expected to experience “coastal squeeze” -- a general narrowing of the extent of the intertidal zone [16,17] and a steepening of the coastal profile [18]. Recent studies provide a disconcerting consensus that sea-level rise may cause substantial habitat loss. Estimates of habitat loss range from 10-27% with 0.30 m of sea-level rise in Scotland [19], to 10% and 57% with 1.0 and 2.0 m of sea-level rise, respectively, in Oregon, USA [20]. Thorner et al. (2014) found that with 0.3-1.0 m of sea-level rise on five rocky headlands in Australia, impacts will be variable but will largely result in substantial habitat loss [21]. Habitat loss is one of the greatest threats to global biodiversity [22,23]. Thus it is critical to evaluate the current state of rocky intertidal ecosystems and assess how sea-level rise may affect these important communities in coming years.

The ecological characteristics that make rocky intertidal systems unique may also make them particularly vulnerable to changes in structure and function as a result of sea-level rise. The rocky intertidal is characterized by patterns of ecological zonation that manifest as distinct bands along the tidal elevation gradient. These bands are generated by a variety of spatially and density-dependent biological mechanisms such as competition [24,25], mutualism [26] and predation [25,27–29], all of which are largely influenced by the physical environment [28,30]. Sea-level rise will cause an upward shift in this banding as the current intertidal is submerged and up-slope habitats become inundated by the sea. While some species may keep pace with this upward shift, many intertidal species are sessile and will likely be incapable of rapidly adjusting their distributions [14]. Additionally it is unclear whether the changing intertidal zone will be suitable for colonization by many species, as habitat characteristics and physical environmental conditions may also change [12]. Thus, coastal squeeze and the rapid upward shift in intertidal area will likely significantly impact the abundance, distribution, and competitive interactions of rocky intertidal species.

### Large-area imaging approach to quantify climate change impacts

The rocky intertidal is one of the most extensively studied ecosystems, and over 75 years of experimentation and monitoring in these habitats has generated an impressive body of fundamental ecological theory and insight into the mechanisms controlling ecosystem structure [24,25,27–29,31–35]. Traditional sampling methods, however, have been largely restricted in their spatial extent, with units of replication on the scale of one to ten square meters. This limits our ability to address ecological processes which operate on larger spatial scales. Determining how climate change will modify landscape scale patterns in biological communities will require an approach that can integrate high-resolution data at ecologically relevant spatial scales. This has been a major technological challenge in the past.

Over the past few years several efforts have advanced the use of innovative geographic information system (GIS) sampling tools and analysis software to provide intensive high-resolution, landscape-scale ecological information in the rocky intertidal [36–39]. Unfortunately, these tools have remained somewhat limited in either spatial coverage or taxonomic resolution due to technological limitations. While remote sensing techniques such as satellite imagery, light detection and ranging (LiDAR), and aerial photography can provide ecological information on broader landscape-scales, they have been generally limited in taxonomic resolution, and researchers have had to rely on traditional field-based methods to provide species identifications. Recent advances in remote sensing, digital imaging, and modern computing now provide researchers new opportunities to explore the interplay between spatial patterns and ecological processes in the rocky intertidal at spatial scales never before possible (from the millimeter to the kilometer scale) [40–42].

Here, we investigated the potential ecological impacts of future sea-level rise on rocky intertidal ecosystems, utilizing a multi-scale approach at two marine reserves in San Diego, CA, USA as a case study. Using a LiDAR dataset, we estimated site-level habitat area changes under a range of sea-level rise scenarios. Using newly available high-resolution large-area imaging tools, we then mapped 720 m^2^ of intertidal habitat and quantified the percent cover, abundance, and density of sessile and mobile organisms at each site. We then used a modelling approach to investigate future sea-level rise driven changes in the cover, abundance, and density of rocky intertidal species. This work takes a critical step toward determining the future impact of sea-level rise on rocky intertidal communities at ecologically relevant scales and provides a novel framework for future monitoring and experimental efforts.

## Methods

### Survey locations and sites

Two rocky intertidal study locations in San Diego County were chosen: the Scripps Coastal Reserve (SCR) and the Cabrillo National Monument (CNM) (Fig 1). These locations are recognized for their ecological and economic importance within the southern California region and are both designated as marine protected areas (MPAs) under California state legislation. CNM sites were studied under a permit granted by the US Department of the Interior National Park Service, Cabrillo (permit #: CABR-2016-SCI-0007), and SCR was studied under a permit granted by the Scripps Coastal Reserve manager (application #: 33783). Long-term ecological monitoring has occurred at one distinct site at SCR (SCR 0) since 1997, and at three sites at CNM (CNM 1-3) since 1990 [43]. All four sites face predominantly west and have a coastal profile and rocky intertidal structure representative of many rocky intertidal coastlines globally [15]. The topography of SCR comprises primarily a large, gently sloping boulder-field backed by steep cliffs, with a large metamorphic dike running south by southwest through the site and providing a distinct upper limit to the intertidal. CNM is composed of a wide, gently-sloping rocky intertidal bench with variable areas of flat sandstone terraces, boulders, scree, and sand accumulation, also backed by steep cliffs [44].

**Figure 1.**
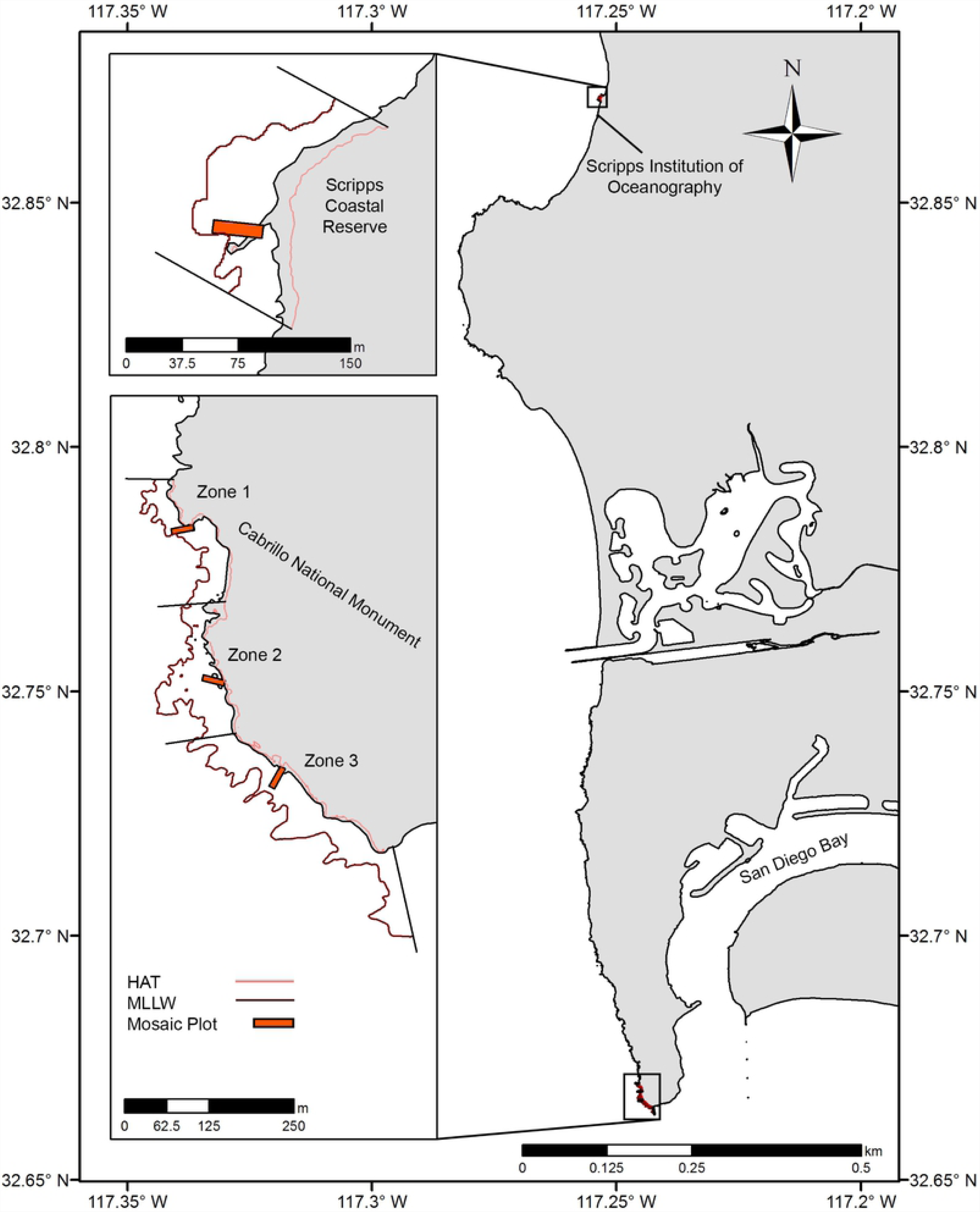
Study site overview. Location of large-area imaging plots (orange rectangles) in San Diego, CA, USA. Sites were selected to fall within long-term monitoring areas (upcoast and downcoast boundaries, black lines), and were bounded by highest astronomical tide (HAT, light red contour) and Mean Lower Low Water (MLLW, dark red contour).

### Sea-level rise scenarios

In its Fifth Assessment Report, the United Nations Intergovernmental Panel on Climate Change (IPCC) projects a rise in global sea-level of between 0.26 m and 0.98 m by 2100 [3]. More recently the National Oceanic and Atmospheric Administration (NOAA) released projections that include sea-level rise extremes of up to 2.5 m on US coastlines [10]. Sea-level rise projections for California specifically generally fall within the range of global projections. Cayan et al. 2008 forecast sea-level rise on the coast of California of 0.11 - 0.72 m by the 2070–2099 period [45]. More recently, the National Research Council (NRC) projected 0.42 - 1.67 m of sea-level rise relative to 2000 levels by 2100 for the coast of California south of Cape Mendocino [5]. For this study, sea-level rise scenarios from 0 – 2.0 m were analyzed in 10.0 cm increments (twenty scenarios) in order to cover the generally accepted potential sea-level rise range for the California region in the next century. By analyzing sea-level rise in increments, we also avoided utilizing projections for specific dates, and thus our analysis is not time specific and is more flexible to the uncertainty of the projections.

### Intertidal area estimation

To estimate rocky intertidal habitat area at survey sites, an open-source LiDAR dataset (the 2009-2011 California Coastal Conservancy Coastal LiDAR Project: Hydro Flattened Bare Earth Digital Elevation Model (DEM)) was used. This dataset was downloaded from the NOAA Office of Coastal Management Data Access Viewer [46]. These data were collected by the California Coastal Conservancy in conjunction with the State of California specifically for shoreline delineation purposes and to inform coastal planning. This dataset has consistent coverage of our target area and a tested data quality of 50.0 cm RMSE horizontal accuracy and 4.8 cm RMSE vertical accuracy [46]. Data was collected with a Leica ALS60 MPiA sensor with 1.0 m nominal post spacing. WGS1984 and Mean Lower Low Water (MLLW) were selected as horizontal and vertical datums during the data download step to allow later embedding of orthophotos and direct referencing to common tidal datums. DEM data was extracted and visualized in ArcMap 10.5 (ESRI 2016. ArcGIS Desktop: Release 10.5 Redlands, CA, USA). Intertidal area was estimated using the DEM Surface Tools Extension for ArcGIS, which provides accurate surface area estimates on a grid by grid basis for raster DEMs using a 3 × 3 cell neighborhood [47].

The area of the intertidal zone was estimated from the DEM constrained within specific boundaries at each survey site. The northern and southern most long-term monitoring plots at each site were selected as fixed upcoast and downcoast boundaries. Tidal datums from NOAA tide stations nearest our survey sites (La Jolla, Station ID: 9410230, and San Diego, Station ID: 9410170 (S1 Table) were selected as vertical constraints that were allowed to migrate upward under each sea-level rise scenario. Mean Lower Low Water (MLLW) and Highest Astronomical Tide (HAT) were chosen as the lower and upper boundaries. Tidal datum elevations are computed from time series of observed tides at NOAA tidal stations. The current National Tidal Datum Epoch (NTDE) is the 1983-2001 period. The DEM was further segmented to specific tidal elevations to allow analyses by intertidal zone (lower, middle, and upper intertidal). MLLW (0 m) was chosen as the lower limit because it is a commonly used datum and because the penetration limitations of LiDAR did not allow for accurate data below the MLLW line. The middle intertidal was designated as the area between mean low water (MLW) and mean high water (MHW). The upper intertidal was designated as the area between MHW and HAT, encompassing the zone that is only covered during the highest high tides, and the spray zone. Long-term monitoring plot locations confirmed the relevancy of our chosen zone designations, as the elevation of plots targeting characteristic lower, middle, and upper intertidal species corresponded closely with our chosen datums (S1 Table). In each scenario, the total area in the intertidal was estimated by adjusting the vertical constraints to the future intertidal extent and summing surface area of all DEM grid cells within the chosen intertidal boundaries.

### Benthic community characterization through fine-scale, large-area imaging

In order to gather fine scale information on biological community composition, one plot was established at each site for image collection and subsequent model creation. Plot locations were selected to encompass all representative vertical zonation within the rocky intertidal. To choose plot locations, ten stratified random coordinates were first generated in ArcMap within the upper intertidal area of the LiDAR DEM covering each survey site. Coordinates were then ground-truthed to ensure they occurred on natural substrate within representative upper-intertidal habitat (evaluated on the presence of representative upper-intertidal benthic species). Of those coordinates falling in appropriate habitat, one was then randomly selected as the upper up-coast corner for each plot. Rectangular 6.0 m × 30.0 m plots were then established running perpendicular to the shoreline along the elevation gradient. This size was determined to be sufficient for accomplishing project goals while balancing image collection and data processing capabilities.

Imagery was collected at the four survey plots between December 2016 and January 2017. Survey dates corresponded with the season’s lowest tides and with associated long-term monitoring surveys. Transect tapes were deployed along predetermined headings to bound the survey plot. A GoPro Hero 5 camera (GoPro Inc., San Mateo, CA) was mounted to a frame on a handheld transect line and passed between two surveyors across the plot every 0.5 m. Images were collected every 0.5 seconds using a linear field of view setting with an equivalent focal length of 24-49 mm. The camera was held approximately 1.0 m above the substrate to maximize overlap of images while also ensuring sufficient image resolution for accurate species identifications. Ten scale bars of known length (0.5 m) were deployed throughout the plot as x, y, z spatial references, and ground control point (GCP) coordinates were collected at the upcoast end of each scale bar. Images were collected over a 9.0 m × 33.0 m area to ensure that the target plot was imaged with sufficient coverage (1.5 m buffer around perimeter) to minimize areas missing data.

### Image processing

Agisoft PhotoScan Professional V.1.3 Structure-from-motion (SFM) software (Agisoft LLC 2014, St. Petersburg, Russia) was used to create 3-dimensional (3D) models, DEMs, and 2-dimensional orthorectified large-area images (e.g. orthophotos) of the intertidal (Fig 2). The details of 3D model and orthophoto creation have been described in detail elsewhere [40–42,48]. Briefly, Agisoft was used to first align imagery and produce a sparse cloud of points extracted from the collected imagery. These sparse clouds were then optimized to correct model geometry and minimize alignment error, and assigned a coordinate system and scale using GCP coordinates and lengths from the scale bars within each plot. Textured dense point clouds were then produced and subsequently meshed to produce fine-scale DEMs of each plot with a 1.0 cm nominal post spacing, providing elevation data for benthic identifications. Finally, top down 2D orthophotos (1.0 mm / pixel resolution) were then produced to allow visual identification and quantification of benthic organisms (Fig 2). This resolution allowed clear identification of all organisms of sizes greater than or equal to approximately 1.0 cm in diameter. Orthophotos and DEMs were exported from Agisoft Photoscan in a raster format and uploaded into ArcMap 10.5. These rasters were then clipped to the targeted extent of each plot, and DEM surface area calculations and benthic identifications were conducted.

**Figure 2.**
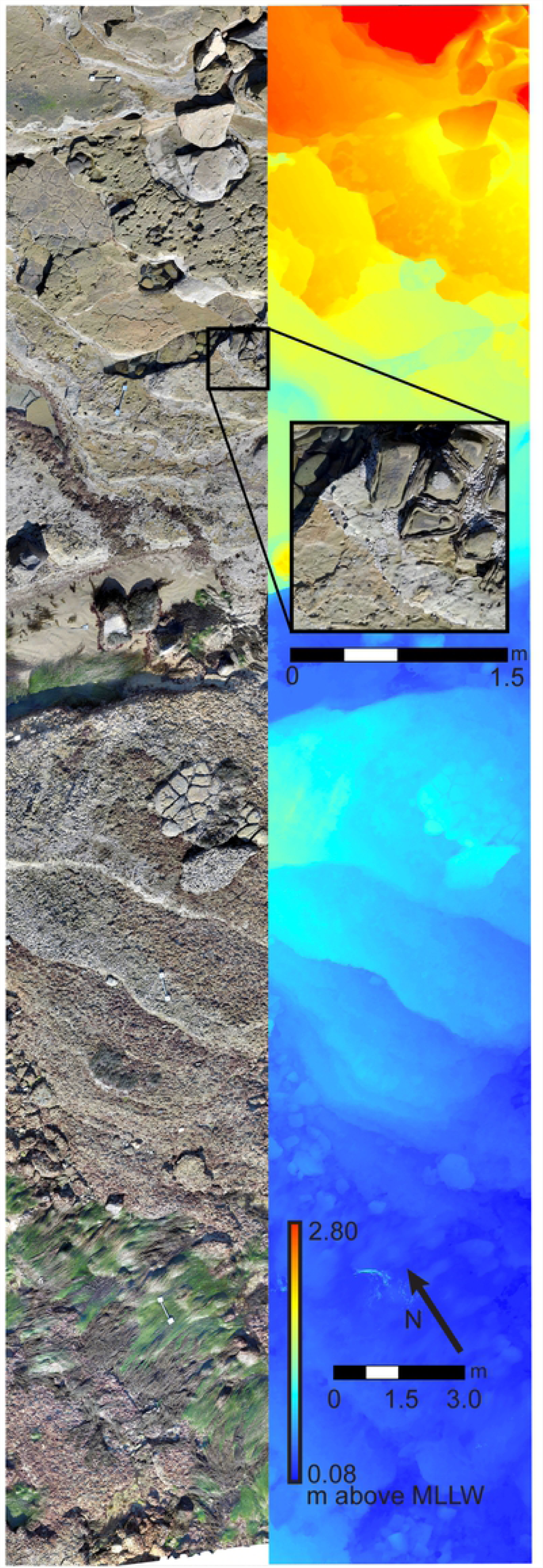
Survey plot photomosaic and Digital Elevation Model (DEM). Example of 6 m × 30 m (180 m^2^) benthic landscape mosaic (left) with zoomed in inset, and DEM (right) for study site Cabrillo National Monument Zone 3.

### Biological data extraction

To determine benthic community composition (% cover) across tidal elevations, stratified random point sampling was conducted within each orthophoto using ArcMap. First, a grid of ninety-six hundred equal sized rectangles (12.5 cm × 15.0 cm) was generated within each plot Ninety-six hundred stratified random points (one within each grid cell) were then generated and these points were manually identified to the highest taxonomic resolution possible (see S2 Table for species identification metadata). Tidal elevation data for each point identification was extracted from the high resolution plot DEMs. This approach allowed quantification of benthic community composition (% cover) across a continuous range of tidal elevation. Benthic community composition was calculated for each species within 10.0 cm elevation bins across each plot using the stratified random point sampling data. Percent cover was calculated by: % 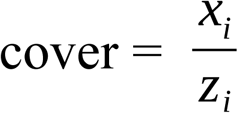, where i = bin number, x_i_ = species stratified random point count in the i^th^ bin, and z_i_ = total stratified random point count for all species within the i^th^ bin.

To provide estimates of densities of targeted long-term monitoring invertebrate species at each site, counts of all invertebrates larger than approximately 1.0 cm in diameter were conducted. This included both motile species (owl limpets (*Lottia gigantea* (Sowerby, 1834)), other limpets (*Lottia spp.)*, black turban snails (*Tegula funebralis* (A. Adams, 1855)), chitons (*Katharina spp.)*, dog whelks (*Nucella spp.*), periwinkles (*Littorina spp.*)) and sessile species (solitary anemones (*Anthopleura sola* (Pearse and Francis, 2000) */ Anthopleura elegantissima* (Brandt, 1835)), pink barnacles (*Tetraclita rubescens* (Darwin, 1854)), gooseneck barnacles (*Pollicipes polymerus* (Sowerby, 1883)), and mussels (*Mytilus californianus* (Conrad, 1837))). Species counts were done at a consistent scale (1:4) to allow accurate identification near the limit of orthophoto resolution. Only invertebrates that could be clearly identified at this scale were counted. density of invertebrates was estimated within the same elevation bins mentioned above across the plots.

Density was calculated by: 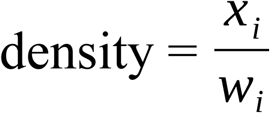, where i = bin number, x_i_ = species point count within the i^th^ bin, and w_i_ = total orthophoto plot surface area within the i^th^ bin.

### Sea-level rise impacts to community composition, abundances, and densities

In order to assess how changes in habitat area as a result of sea-level rise may influence the rocky intertidal community, the cover of benthic organisms (area, m^2^), abundance, and density of invertebrates was estimated for each site under each sea-level rise scenario.

#### Species benthic area cover

To estimate how sea-level rise will change benthic community composition, the total area (m^2^) covered by each species in each scenario at each site was calculated as: 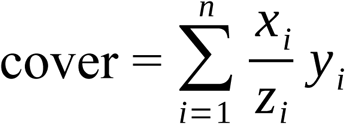, where i = bin number, x_i_ = species stratified random point count within the i^th^ bin, z_i_ = total point count for all species within the i^th^ bin, and y_i_ = site surface area within the i^th^ bin.

#### Scenario abundance estimates

To estimate how sea-level rise will alter invertebrate populations, the abundance (# individuals) for each species in each scenario at each site was calculated as: 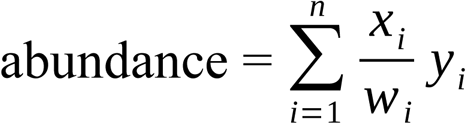, where i = bin number, x_i_ = species point count within the i^th^ elevation bin, w_i_ = sampled orthophoto plot surface area within the i^th^ elevation bin, and y_i_ = landscape surface area within the i^th^ bin.

#### Scenario density estimates

In order to estimate overall density for each species in each scenario at each site the above abundance estimate values were divided by the total intertidal area within each site in each scenario. Density (# / m^2^) was calculated as: 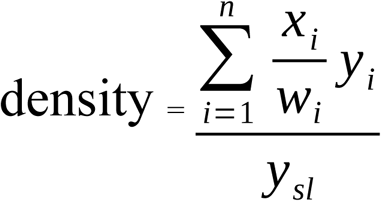, where s = scenario number, l = site, i = bin number, x_i_ = species point count within the i^th^ elevation bin, w_i_ = sampled orthophoto plot surface area within the i^th^ elevation bin, y_i_ = landscape surface area within the i^th^ bin, and y_sl_ = landscape surface area across the entire intertidal for scenario s at site l.

## Results

### Change in intertidal habitat area and zonation with sea-level rise

Using a LiDAR elevation dataset, we estimated habitat area within current and future intertidal elevation ranges at each of our study sites under scenarios of 0 - 2.0 m of sea-level rise and found that sea-level rise will significantly reduce total intertidal habitat area (m^2^) (Fig 3). Following a sea-level rise trajectory consistent with the observed trend in San Diego, CA, (approximately 20.0 cm by 2100), total intertidal habitat area loss will be on average 29.88 % (± 3.78, SE) across study sites. Under the IPCC upper-end global projection of 1.0 m by 2100 [3], this value will reach 77.72% (± 4.65). Under the NRC upper-end projection for California [5] of 1.7 m, this value will rise to 85.32% (± 2.33). Habitat loss will be greatest for the lower and middle intertidal zones, which currently occupy a broad intertidal shelf that will rapidly become subtidal as sea-levels rise (Fig 3). Under scenarios greater than 0.2 m the lower intertidal will nearly always experience the greatest proportional habitat area loss, followed by the middle, then upper zones (Fig 3). As a result, we expect that the proportional contribution of each zone to total intertidal area will shift, with the contribution of the lower intertidal diminishing, and that of the middle and upper zones increasing (Fig 3, S1 Fig).

**Figure 3.**
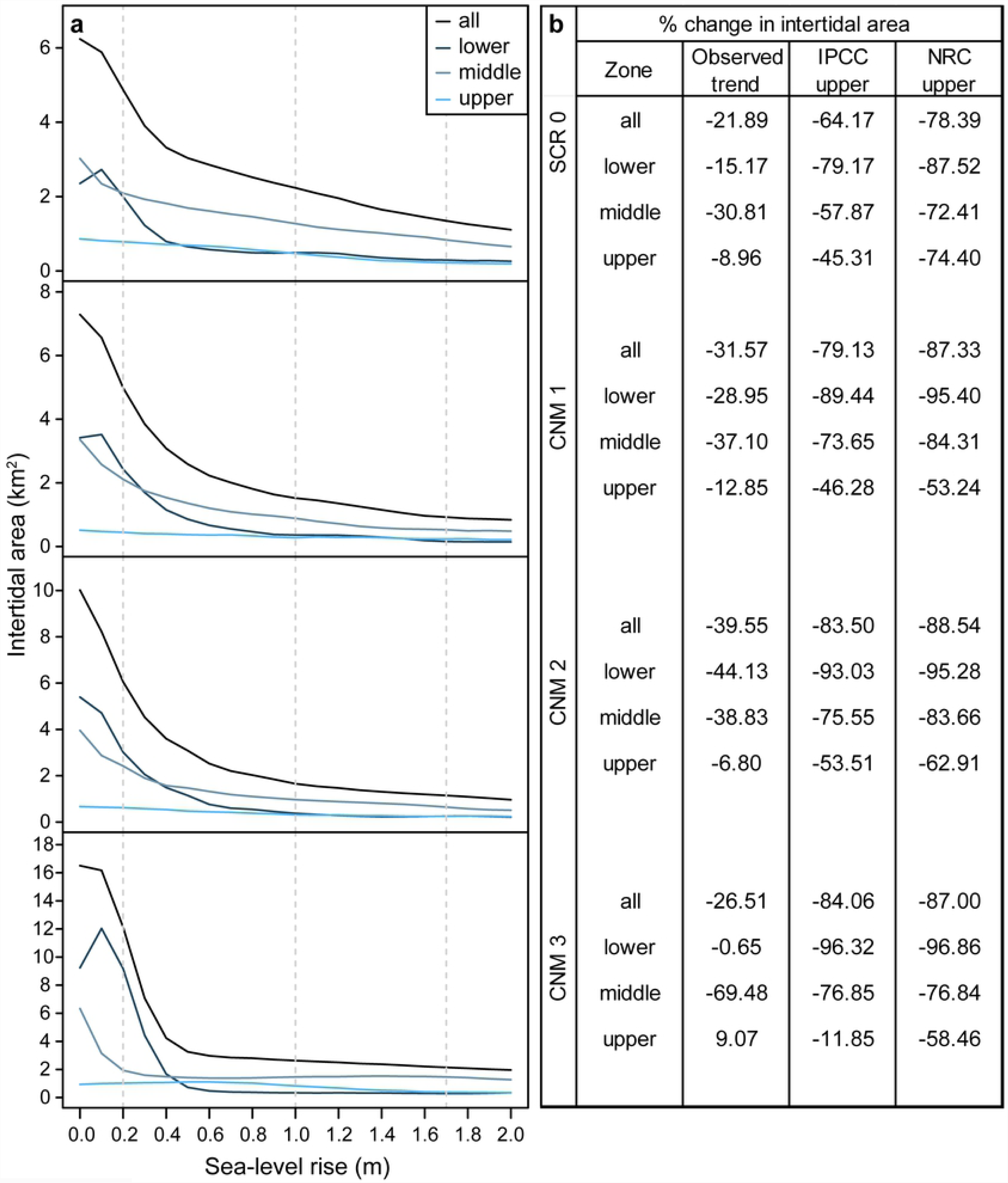
Intertidal area changes with sea-level rise. a. site and zone level intertidal habitat area under 0 – 2.0 m of sea-level rise at four survey sites. b. site and zone level intertidal area change (% change) under three sea-level rise scenarios. Both panels show significant habitat area loss at study sites.

### Impacts to benthic community composition with sea-level rise

With little exception, we found that sea-level rise will result in lower overall habitat area and thus benthic area cover (m^2^) of rocky intertidal organisms. Importantly, species will experience benthic cover changes of different magnitudes, resulting in changes in community structure as the relative abundance of different species shifts (Fig 4). The most pronounced changes will occur in the first half meter of sea-level rise for all species. At 0.5 m of sea-level rise, we estimate a mean decrease in total benthic cover across all species and study sites of 56.95% (± 2.40). Benthic cover changes will be most pronounced for those taxa that primarily occupy lower and middle intertidal habitats, such as articulated coralline algae, brown algae, red foliose algae, and turf algae species, and surfgrass (see S2 Table for species identifications and classifications). For example, we estimate that cover of articulated coralline algae will decrease by an average of 83.74% (± 4.72 SE; range 70.10 - 91.77) across our study sites under the IPCC upper-end projection. Under the more extreme NRC projection, we expect decreases of cover of up to 98.22%, 97.42% and 97.20% for chainbladder kelp (*Stephanocystis setchellii* ((N.L.Gardner) Draisma, Ballesteros, F.Rousseau & T.Thibaut, 2010: 1340)), wireweed (*Sargassum muticum* ((Yendo) Fensholt, 1995), and surfgrass (*Phylospadix spp.*), respectively. Benthic cover is expected to change less dramatically for taxa occupying primarily upper intertidal habitat, such as mussels, barnacles, crustose coralline algae, and green algal turf. For example, we expect that cover of barnacles (*Balanus/Cthamalus spp.*) will decrease by an average of 52.99 % (± 12.01; range 20.38 - 77.84) across our study sites under the IPCC upper-end projection.

**Figure 4.**
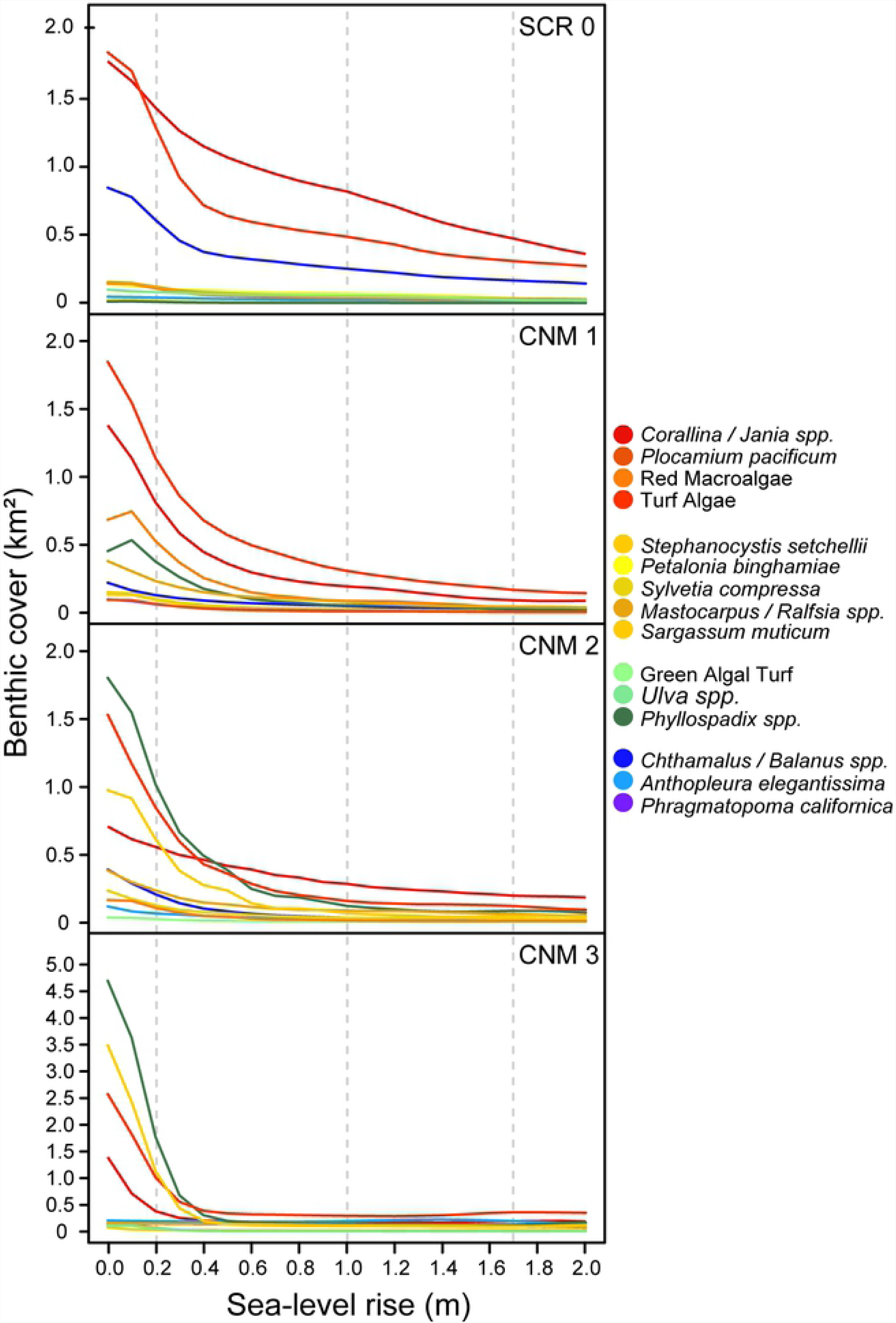
Sea-level rise impacts to benthic community cover. Estimated current and future intertidal area coverage for 10 most common benthic community members at each survey site under 0 – 2.0 m of sea-level rise, showing decreases for all benthic space occupiers.

### Invertebrate abundance and density estimates for sea-level rise scenarios

We estimate nearly ubiquitous declines in numerical abundance of sessile and mobile invertebrates with sea-level rise (Fig 5). Lower and middle intertidal taxa will exhibit greater population declines than upper intertidal taxa, and the largest declines will be observed in the first 0.5 to 1.0 meter of sea-level rise. For example, our results suggest that the abundance of green solitary anemone (*Anthopleura sola/xanthogrammica*) will decrease by an average of 64.37% (± 8.66) and 76.20% (± 15.60) across study sites at 0.5 and 1.0 m of sea-level rise, respectively. In contrast, we estimate smaller declines in the abundance of upper intertidal periwinkles (*Littorina spp.*) with 26.22% (± 14.9) and 51.19% (± 0.17) and upper intertidal goose barnacles with 29.10% (± 5.60) and 47.14% (± 7.54) across study sites at 0.5 and 1.0 m of sea-level rise, respectively. Across all intertidal invertebrate taxa identified and sites, our results suggest overall mean decreases in abundance of 55.82% (± 4.25) and 66.92% (± 4.09) under the IPCC and NRC projections, respectively.

**Figure 5.**
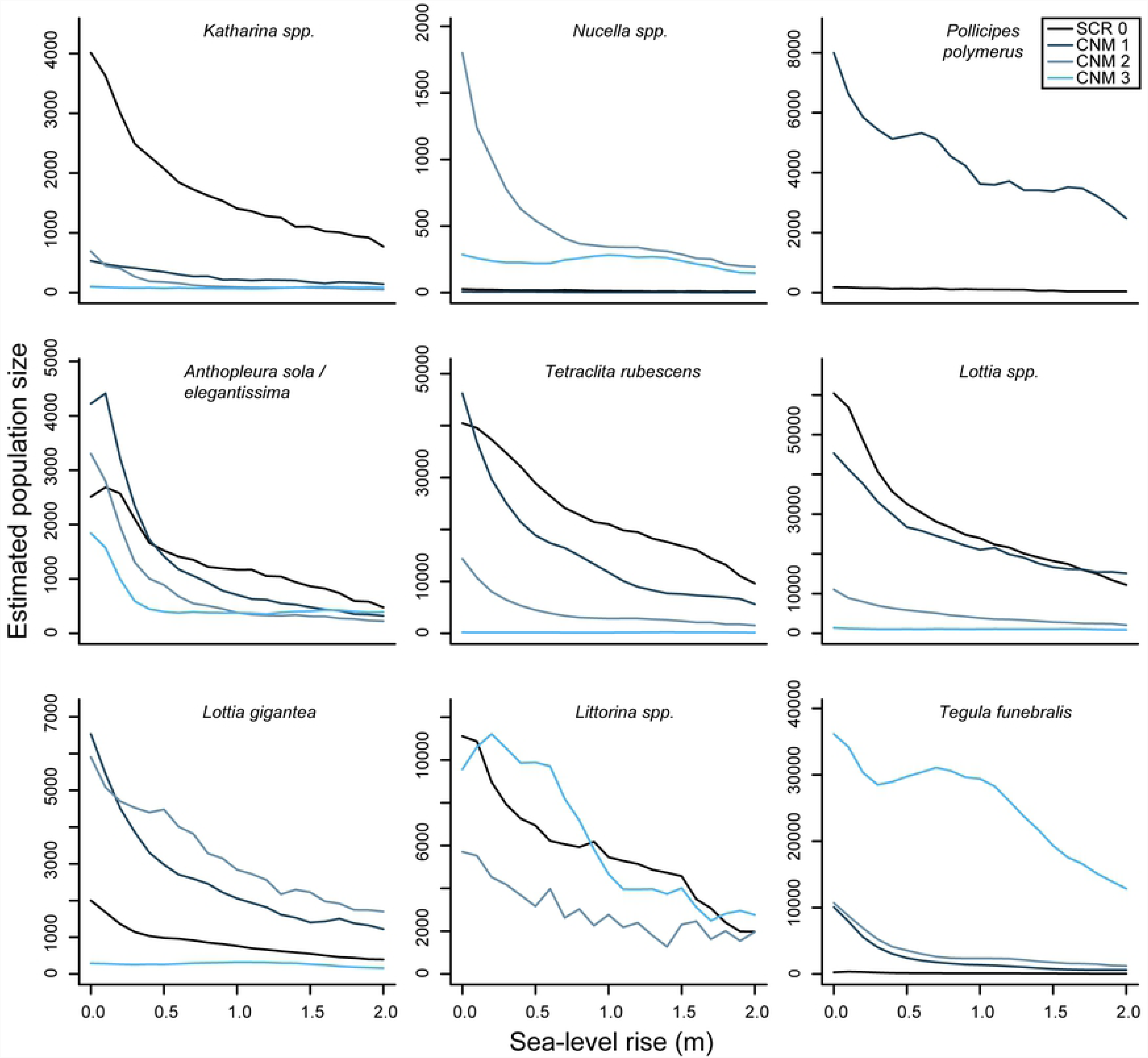
Sea-level rise impacts to rocky intertidal invertebrate abundances. Estimated current and future abundances for large invertebrates at survey sites under 0 - 2.0 m sea-level rise.

Despite expected declines in overall abundance for many species, we expect to see an increase in density resulting from the relatively large predicted declines in habitat area under most sea-level rise scenarios (Fig 6). Nearly all species exhibit significant increases in overall density in the first 0.5 m of sea-level rise at one or more sites. For example, our results suggest an average increase in density of 85.35% (± 64.57) for lower intertidal chitons and of 177.65% (± 107.71) for upper intertidal periwinkles (*Littorina spp.*) across sites under 0.5 m of sea-level rise. Beyond 0.5 m of sea-level rise the predicted density trends are more variable, though densities generally increase with continued loss of colonizable habitat. The major exception was seen with the green anemone (*Anthopleura sola/xanthogrammica*) at CNM 1 and CNM 2, which we predict will generally decrease in density as a result of the large estimated declines in population size at these sites.

**Figure 6.**
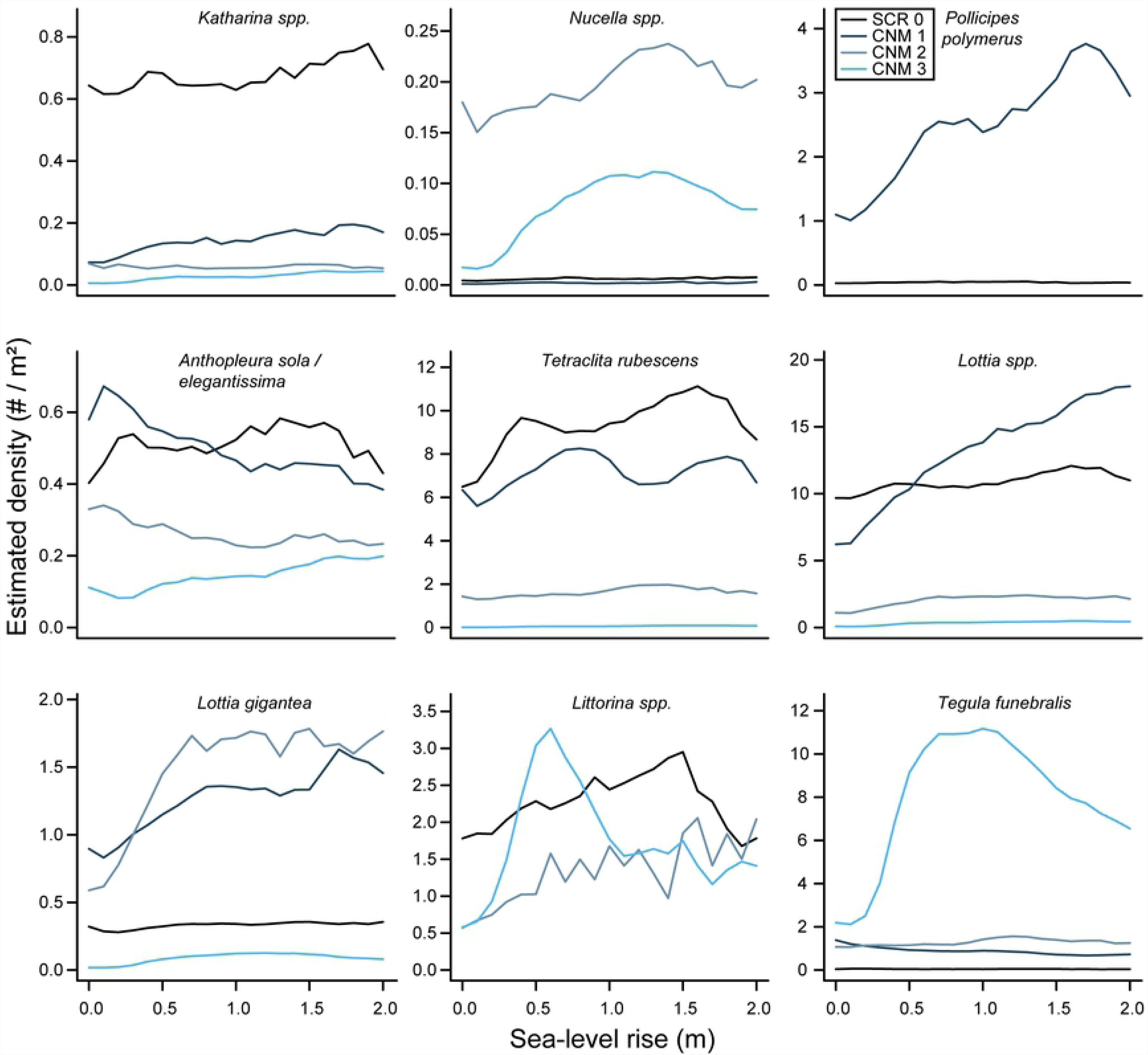
Sea-level rise impacts to rocky intertidal invertebrate densities. Estimated current and future invertebrate densities at study sites.

## Discussion

Our results suggest that sea-level rise will significantly reduce total rocky intertidal habitat area at our study sites. Changes will neither occur uniformly across time nor space, but rather will be most pronounced during the first meter of sea-level rise and within the lower and middle intertidal zones. As seas rise, the broad intertidal bench present at our study locations will quickly be submerged, resulting in a first-rapid, then more gradual loss of intertidal habitat area. We also model variable but generally negative impacts to the rocky intertidal communities at our survey sites, even under modest sea-level rise scenarios well within the range projected for the next century. As the amount of available habitat decreases we predict reductions in benthic cover for all major benthic space occupiers. We also predict significant population declines for invertebrate species. Finally, despite declines in overall abundance, we predict that densities of invertebrates will generally increase as habitat area decreases.

### Impacts to rocky intertidal community structure

Habitat area is a limiting factor affecting growth and abundance of rocky intertidal species [32]. Thus, the changes in habitat area estimated here will likely significantly impact community structure. As habitat area is compressed, biotic interactions known to drive community structure will likely change. Both competition and predation will likely intensify as densities of sessile and mobile invertebrates increase. Additionally, interactions that were otherwise rare or non-existent due to the spatial separation between species may become intensified or be created as species are spatially compressed. Further, because rocky intertidal organisms exhibit distinct distributions across tidal elevation and a range of life history strategies, the impact of sea-level rise will likely be non-uniform across the rocky intertidal community both taxonomically and through time. Our results suggest that lower and middle intertidal species will generally experience the greatest losses of benthic cover and abundance. Because these species will also experience extensive habitat loss more quickly than upper intertidal species, it is possible that they will struggle to compete for space as they are forced to move upward. If dominant upper intertidal benthic space occupiers, such as the California Mussel (*Mytilus californianus*), experience little habitat loss under early sea-level rise scenarios and have little competition for space in the largely bare substrate above their current distributions, they may gain a distinct competitive advantage. Long-lived species with high habitat affinities and small home ranges, such as owl limpets (*Lottia gigantea* (Sowerby, 1834)) and the critically endangered intertidal black abalone (*Haliotis cracherodii* (Leach, 1814)) will potentially be more heavily impacted than short-lived species with low habitat affinities, such as many barnacles, even if these species occupy the same tidal zone. These species may experience significant change within their lifespans and are more likely to experience loss of necessary microhabitats as conditions change due to their narrow habitat requirements.

### Impacts to rocky intertidal function

As sea-level rise changes species interaction networks and competitive hierarchies, the function of the rocky intertidal ecosystem will also potentially shift. Our results suggest that as sea-levels rise, the middle and upper intertidal will occupy a greater proportion of total intertidal area (S1 Fig). Lower shore habitats are generally recognized to be richer in terms of species diversity and higher in productivity [49], thus declines in this zone may be of particular consequence. Loss of primary producers common in the lower intertidal may drive a subsequent reduction in herbivores in this region and a reduction in nutrient and biomass export to adjacent areas [50]. Additionally, loss of pools common in the lower intertidal as a result of a general steepening of the coastal profile may decrease important nursery habitat for offshore species such as the opaleye (*Girella nigricans*(Ayres, 1860)) [51], and foraging habitat for commercially important species such as the spiny lobster (*Panulirus interruptus* (Randall, 1840)) [52] as well as many seabirds [20]. As seas rise it will be crucial to examine the impacts of such shifts in community structure to ecosystem function to inform management policies.

### Future directions

Our study provides a framework to evaluate climate change impacts on one the world’s most important marine ecosystems at a scale not previously possible. We found compelling evidence that sea-level rise will significantly and profoundly affect species inhabiting this habitat. Future studies can improve upon this approach by incorporating additional information on physical parameters that are known to influence spatial heterogeneity in rocky intertidal community organization and that are likely to evolve under global climate change, such as temperature [53], ocean chemistry [54,55], and wave intensity [45]. Further coverage of much larger areas of coastline will also allow researchers to more precisely predict the future impacts of climate change on the rocky intertidal.

The rocky intertidal is the most accessible of marine environments, and is of immense recreational, commercial, and educational value to coastal societies worldwide. These systems are likely to be substantially modified by large-magnitude global sea-level rise on an accelerated and uncertain timeline within the next century. The implications of our results are wide reaching, highlighting the need for ecosystem-scale evaluations in order to quantify and visualize the global change impacts that will modify the structure and function of this unique ecosystem. Similar approaches are needed more broadly for global coastlines in order to understand how to manage and mitigate impending global change impacts [42].

## Acknowledgements

The authors would like to acknowledge C. Amir, L. Bonito, D. Chargualaf, A. Martinez for their assistance collecting field data, J. Jones and K. Lombardo for assistance with acquiring permits and accessing Cabrillo National Monument sites, I. Kaye for providing permits to access the Scripps Coastal Reserve, J. Jenness for advice on analyzing DEM data in GIS and on the use of the DEM Surface Analyst extension for ArcGIS, S.A. Sandin for modeling the use of large area imaging for studying spatial ecology, E. Parnell for his natural history guidance, V. Petrovic for his assistance with 3D model visualization, and N. Pederson for assistance with image processing.

## Supporting information captions

**S1 Table. Site Tidal datums from NOAA tidal stations.** Tidal datums were used to segment the intertidal into zones (upper, middle, lower). Data presented in m in reference to MLLW.

**S2 Table. Species Identification Metadata.** Table shows species and functional group level identifications used for stratified random point counts and invertebrate counts.

**S1 Figure.** Proportion of intertidal area contributed by each tidal zone under 0 – 2.0 m of sea-level rise for four survey sites.

**S1 Dataset. Intertidal area estimate data.** Estimates of intertidal area and percent change in intertidal area (site- and zone-level) for sea-level rise scenarios from LiDAR raster DEM.

**S2 Dataset. Species benthic cover estimate data.** Site-level estimates of benthic area cover and percent change in benthic area cover of benthic community members for sea-level rise scenarios.

**S3 Dataset. Abundance estimate data.** Site-level estimates of abundance of invertebrate taxa for sea-level rise scenarios.

**S4 Dataset. Density estimate data.** Site-level estimates of density of invertebrate taxa for sea-level rise scenarios.

